# Structural organization and energy storage in crosslinked actin-assemblies

**DOI:** 10.1101/237412

**Authors:** Rui Ma, Julien Berro

## Abstract

During clathrin-mediated endocytosis in yeast cells, short actin filaments (< 200nm) and crosslinking protein fimbrin assemble to drive the internalization of the plasma membrane. However, the organization of the actin meshwork during endocytosis remains largely unknown. In addition, only a small fraction of the force necessary to elongate and pinch off vesicles can be accounted for by actin polymerization alone. In this paper, we used mathematical modeling to study the self-organization of rigid actin filaments in the presence of elastic crosslinkers in conditions relevant to endocytosis. We found that actin filaments condense into either a disordered meshwork or an ordered bundle depending on filament length and the mechanical and kinetical properties of the crosslinkers. Our simulations also demonstrated that these nanometer-scale actin structures can store a large amount of elastic energy within the crosslinkers (up to 10*k*_B_*T* per crosslinker). This conversion of binding energy into elastic energy is the consequence of geometric constraints created by the helical pitch of the actin filaments, which results in frustrated configurations of crosslinkers attached to filaments. We propose that this stored elastic energy can be used at a later time in the endocytic process. As a proof of principle, we presented a simple mechanism for sustained torque production by ordered detachment of crosslinkers from a pair of parallel filaments.

## I. INTRODUCTION

The cytoskeleton protein actins assemble into three major structures in yeast cells, including endocytic actin patches, actin cables, and the contractile ring [1, 2]. In actin cables and the contractile ring, formin-nucleated actin filaments are crosslinked into long bundles with a length on the order of microns [3–5]. Computational models of these actin structures typically treat actin filaments as semi-flexible polymers that are connected by rigid segments [6–9]. In contrast, the organization of the actin network in actin patches formed during clathrin-mediated endocytosis is drastically different from that in actin cables or the contractile ring. The length of filaments in actin patches is strongly limited by capping and severing proteins [10], and mathematical modeling predicted that the average length of filaments is less than 200 nm [11]. Filaments of this length scale can be considered as straight rods, because the persistence length of actin filaments is on the order of 10*μ*m [12–14], which allows them to sustain forces larger than 10pN without buckling [15].

At the endocytic actin patch, a small area of the flat plasma membrane invaginates towards the cytoplasm upon assembly of actin. In budding yeast, the invagination elongates up to 140nm in depth, and then is pinched off, releasing a tear-shaped vesicle [16]. Actin is essential for many of these steps, from the initiation of invagination to vesicle scission [17–19]. Despite extensive experimental work that characterized the overall dynamics of assembly, disassembly and ensemble movements of proteins of the actin meshwork [19–26], the precise structural organization of actin filaments within the endocytic patch remains unknown. Indeed, individual filaments are not resolvable even in electron micrographs, in which the actin network appear as a ribosome-exclusion zone, which is about 200nm in depth and 100nm in width [16].

Actin crosslinking proteins play a crucial role in determining the mechanical responses of the actin network to force perturbation [27–29]. Fimbrin (Fim1p) is the second most abundant protein recruited to the endocytic patch during clathrin-mediated endocytosis in fission yeast, after actin [20]. It has two actin binding domains that enable it to crosslink adjacent filaments. Deletion of fimbrin results in significant defects in endocytic internalization [24, 30, 31]. *In vitro* experiments have shown that fimbrin efficiently bundles long actin filaments, but bundling efficiency is reduced in the presence of capping protein as a result of decreased filament length [31]. It remains unclear how this length-dependent bundling activity arises and how this activity is related to the role of fimbrin during clathrin mediated endocytosis.

Internalization of the endocytic membrane is hindered by the high turgor pressure (*P* ~ 0.8 × 10^6^Pa [32, 33]) inside yeast cells [34]. Under such high pressure, theoretical studies suggest that the force needed to initiate membrane invagination is on the order of 3000pN [35, 36] and actin polymerization is thought to provide the driving force. However, assuming no more than 150 filaments are simultaneously generating the force [11, 20, 21], each of these filaments has to generate a force of at least 20pN, an order of magnitude larger than the maximum polymerization force of ~ 1pN of actin filaments measured *in vitro* [37]. This number of 20pN is likely an underestimate since the calculation here uses a very generous estimate for the number of growing filaments, up to 20-fold of what mathematical modeling predicts [11]. Therefore, actin polymerization alone is not enough to provide the force necessary to elongate a clathrin-coated pit. Even though type-I myosins participate in endocytosis, their low power output over a narrow range of forces suggest that they are more likely force-sensing tethers rather than force generators [38–40].

In this paper, we present a computational model for dynamic crosslinking of rigid actin filaments in conditions relevant to clathrin-mediated endocytosis. We show that kinetic and mechanical properties of the crosslinkers finely tune the structural transition of actin network between bundles and meshworks. In addition, we show that the chemical binding energy is converted into elastic energy upon binding of crosslinkers. The elastic energy stored in individual crosslinkers is significantly higher than their thermal energy. This surprising property is a consequence of the helical pitch of actin filaments, which leads to torsional strains between crosslinkers attached to a common pair of filaments. We discuss the mechanical implications of these torsionally stressed crosslinkers and propose a possible mechanism to generate directed rotation of filaments by orderly detaching the crosslinkers.

## II. MODEL OF CROSSLINKED RIGID ACTIN FILAMENTS

We model actin filaments as rigid cylindrical rods with subunits that carry a helical pitch (Fig. 1 A). The length of filaments is restricted to the range of actin filament size during endocytosis (81nm to 216nm), around 2 orders of magnitude shorter than their persistence length ~ 10μm. Filaments of this length scale remain virtually straight under ~ 10pN of force and untwisted under ~ 100pN · nm of torque (see Appx. E). Therefore we can neglect bending and twisting, and describe the motion of a filament by its translational velocity **V**_*c*_ of center of mass, and angular velocity **Ω** relative to the center of mass. Details of the model can be found in Appx. A.

**FIG. 1.**
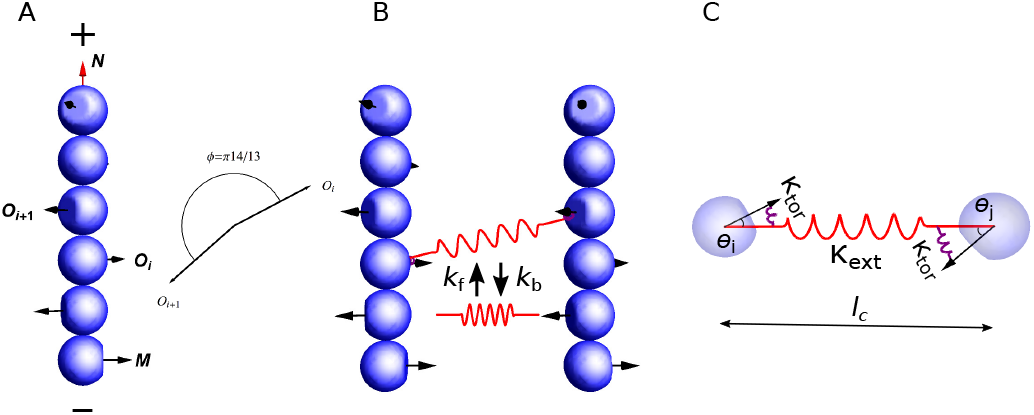
Model description. (A) Actin filaments are described as rigid rods made of subunits carrying an orientational vector **O**_*i*_ that represents the normal vector to the binding surface for an actin crosslinker. The orientation of a filament is described by the unit vector N, pointing from the pointed end (−) to the barbed end (+), and the first subunit’s orientational vector **M** = **O**_1_. Two consecutive subunits have an angle of *π*14/13 in their orientations. (B) Crosslinker turnover is described by stochastic formation and breakage of bonds between two actin subunits in different filaments, with rate constants *k_f_* and *k_b_*, respectively. (C) Each crosslinker is modeled as a combination of springs with extensional stiffness *κ*_ext_, which characterizes the flexibility of the stretchiness *l_c_* between the two actin binding domains, and torsional stiffness *κ*_tor_, which characterizes the flexibility of the angles *θ_i_* and *θ_j_* between the crosslinker and the two actin subunits it is bound to.

We model actin crosslinkers as elastic springs that connect two actin subunits in different filaments. This elasticity is the simplest model to account for: (i) the intra-molecular flexibility between both actin binding domains, and (ii) the flexibility in the binding interface between each actin binding domains and the actin subunits they are bound to. Specifically, the elastic energy *E* of a crosslinker is composed of an extensional part *E*_ext_ and a torsional part *E*_tor_, *E* = *E*_ext_ + *E*_tor_. The extensional energy accounts for (i), (ii),

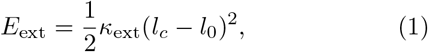

where *κ*_ext_ denotes the extensional stiffness, *l_c_* denotes the length, and *l*_0_ denotes the rest length of the crosslinker (Fig. 1 C). The torsional energy accounts for

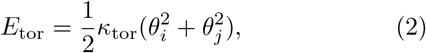

where *κ*_tor_ denotes the torsional stiffness, *θ_i_* and *θ_j_* denote the angles between the actin subunits and the axis of the crosslinker (Fig. 1 C). Note that this torsional spring has no direct influence on the rotation of one filament around the axis of the crosslinker, e.g. it does not capture any preference for parallel or anti-parallel bundles. No experimental values for the extensional and torsional stiffnesses of fimbrin are available in the literature. For our simulations, we used values within a few orders of magnitude of stiffnesses measured for fascin and antibodies [41, 42]. Crosslinker turnover is modeled as Poisson processes with a crosslinker formation rate constant *k_f_* and a breakage rate constant *k_b_*, which increases exponentially with the total elastic energy (Fig. 1 B). A more detailed description is presented in Appx. B.

The length of filaments is kept constant in a given simulation and the total number of actin subunits in the filaments is fixed to *N*_actin_ = 7000 for all simulations [20]. The maximum occupancy of crosslinkers on filaments is constrained to remain under 25% (or 1 crosslinker for 4 subunits), which is equivalent to a maximum of 875 attached crosslinkers, close to the peak value ~ 900 measured experimentally [20]. We initiate each simulation with uncrosslinked filaments that are randomly positioned and oriented. Reflecting boundary conditions are imposed to enforce filaments stay in a cubic box of 500nm in size. Simulations are performed using the reference values listed in Table I unless otherwise mentioned.

**TABLE I.** List of parameters

| Symbol | Meaning | Reference value | Varied Range | Unit |
| --- | --- | --- | --- | --- |
| $L$ | Filament length | 135 | 81-216 | nm |
| $b$ | Filament diameter | 6 | 6 | nm |
| $\delta$ | Length elongation of filaments by incorporation of a subunit | 2.7 | 2.7 | nm |
| $\kappa_{\text{ext}}$ | Extensional stiffness of the crosslinker | 0.1 | 0.01-10 | pN/nm |
| $\kappa_{\text{tor}}$ | Torsional stiffness of the crosslinker | 10 | 0.1-100 | pN · nm · rad<sup>-1</sup> |
| $l_0$ | Rest length of the crosslinker | 10 | 10 | nm |
| $k_f$ | Formation rate constant of the crosslinker | 1 | 0.01-1 | s<sup>-1</sup> |
| $k_b^0$ | Strain-free breakage rate constant of the crosslinker | 10 | 0.01-100 | s<sup>-1</sup> |
| $E_c$ | Critical energy of the crosslinker | 10 | 10 | $k_B T$ |
| $r_c$ | Reaction distance between actin subunits for crosslinker binding | 20 | 20 | nm |
| $f_{\text{st}}$ | Magnitude of the repulsive force between filaments | 100 | 100 | pN |
| $k_B T$ | Thermal energy unit at room temperature | 4.1 | 4.1 | pN · nm |
| $N_{\text{actin}}$ | Total number of actin subunits | 7000 | 7000 | |
| $\eta$ | Viscosity of the medium | 10 | 0.1-10 | Pa · s |
| $\Delta t$ | Simulation time step | 10<sup>-5</sup> | 10<sup>-7</sup> – 10<sup>-5</sup> | s |

To quantify the structure of actin filaments, we introduced the global and local nematic order parameters *S*_global_ and S_local_. These quantities characterize the degree of alignment between filaments in the entire simulation space, or in a local neighborhood, respectively. Their values range from 0 to 1, and a larger value indicates that filaments are more aligned with each other globally, for *S*_global_, or locally, for *S*_local_. Note that, in practice, the minimum reachable value for *S*_local_ is usually close to 0.4 (Fig. S6 [43]). The detailed mathematical definition of these parameters can be found in Appx. D.

## III. RESULTS

### A. Long filaments form bundles and short filaments form meshworks

We first studied how filament length influences the structure of actin clusters. For crosslinking rates *k*_f_ below 0.1*s*^−1^, the number of attached crosslinkers remained small, and actin filaments did not organize into higher order assemblies (Fig. S1 [43]). When the crosslinking rate *k*_f_ was high enough, initially disconnected short filaments (81nm) quickly formed small clusters that eventually coalesced into three to four larger clusters (Fig. 2 A). The number of attached crosslinkers rapidly saturated to a dynamic steady-state (Fig. 2 C), as the crosslinkers underwent constant turnover. Filaments within the cluster are organized into a disordered meshwork, which is characterized by a small local nematic order parameter (*S*_local_ ~ 0.4, Fig. 2 B). Long filaments (216nm) rapidly aligned with their neighbors into small bundles (Fig. 2 D), as indicated by the fast convergence of the local nematic order parameter *S*_local_ to its steady state value around 1 (Fig. 2 E). Throughout the simulation, these locally aligned filaments remained connected with each other, and the number of clusters remained small (Fig. 2 F). These connected bundles slowly adjusted their orientations to eventually coalesce into a few large bundles.

**FIG. 2.**
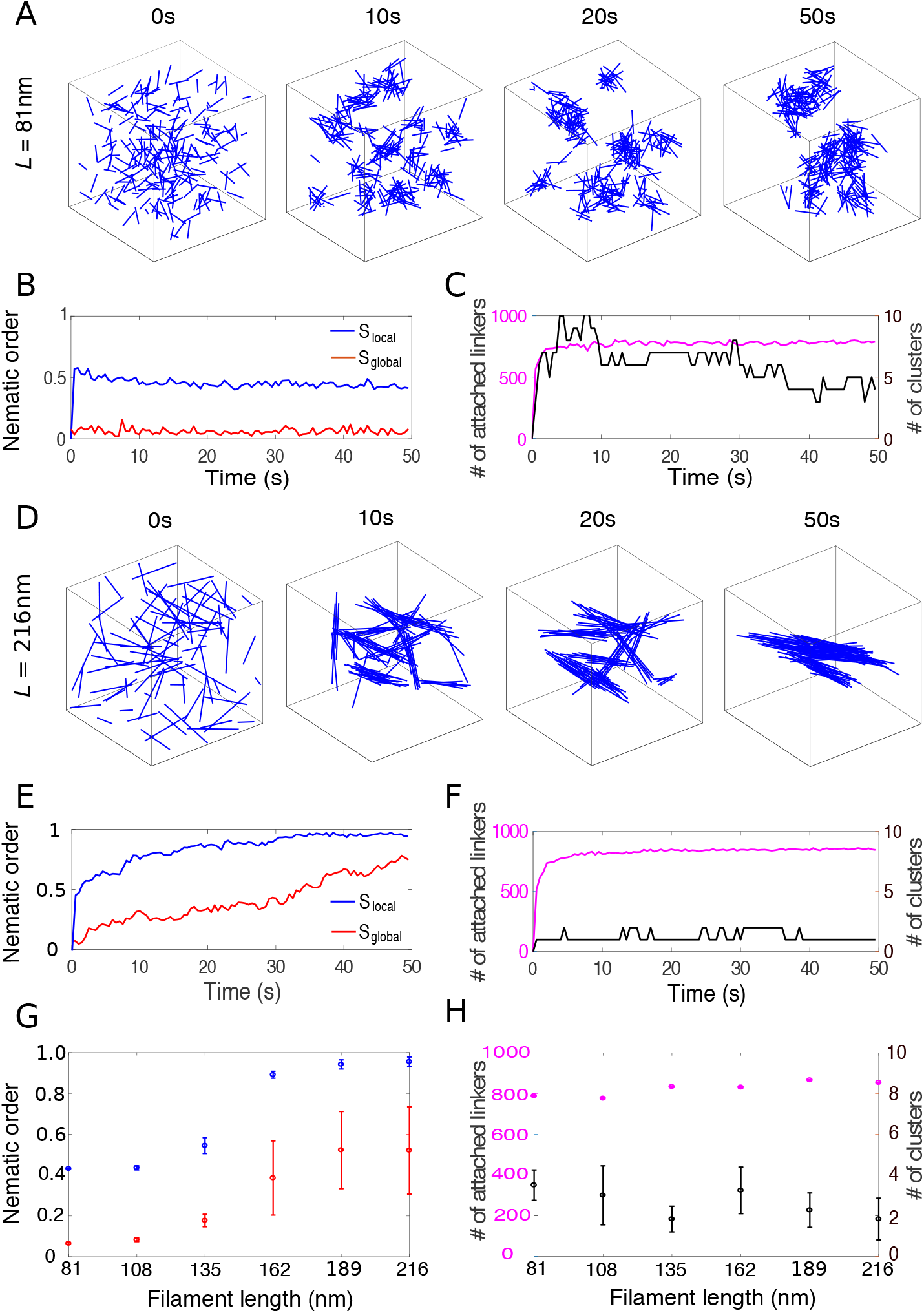
The structure of crosslinked actin networks depends on filament length. (A) Snapshots of the actin network for filaments of 81nm in a cubic box of 500nm in size. Each filament is represented by a blue line. Crosslinkers are not shown for clarity. (B) Local (blue) and global (red) nematic order parameters of the actin network over the course of the simulation shown in (A). (C) Number of attached crosslinkers (magenta, left axis) and the number of clusters (black, right axis) for the simulation in (A). (D-F) Similar figures as in (A-C) but for 216nm-long filaments. (G) Local (blue) and global (red) nematic order parameters as a function of filament length. (H) Number of attached crosslinkers (magenta, left axis) and number of clusters (black, right axis) as a function of filament length. In (G) and (H), for each simulation, the means of the metrics were calculated from the data between 40s to 50s and the error bars indicate standard deviation over 10 simulations.

When filament length was increased from 135nm to 162nm, the structure of the actin network transitioned from meshwork to bundle, indicated by the sharp increase of the local nematic order parameter *S*_local_ from ~ 0.5 to above 0.9 (Fig. 2 G). The global nematic order parameter *S*_global_ showed similar trend as *S*_local_, but with a smaller magnitude, since filaments formed 2 to 4 independent clusters (Fig. 2 H). Altogether these results show that crosslinked actin filaments with a size and crosslinker density comparable to what is measured during endocytosis can self-organize into either meshworks (for short filaments) or bundles (for long filaments), and the phase transition is tightly controlled by filament length.

### B. Crosslinkers with high extensional stiffness drive bundle formation

Next, we explored the influence of the mechanical properties of actin crosslinking proteins on the organization of actin filaments. The organization of medium length filaments (135nm) varied dramatically for different combinations of of *κ*_ext_ and *κ*_tor_ (Fig. 3 A-C). For low extensional stiffness *κ*_ext_, actin filaments organized into a meshwork (Fig. 3 A), while they formed bundles for higher *κ*_ext_ values (Fig. 3 B). The transition between meshwork and bundle was tightly controlled, since the local nematic order parameter had a sharp increase around *κ*_ext_ = 0.1pN/nm (Fig. 3 E, red) at given *κ*_tor_ = 10pN · nm · rad^−1^. This sharp transition was even more pronounced for longer filaments (189nm), as reported in both local and global nematic order parameters (Fig. 3 E and Fig. S2 A, orange [43]). In contrast, when filaments were short (81nm), the transition was relatively smooth (Fig. 3 E, blue).

**FIG. 3.**
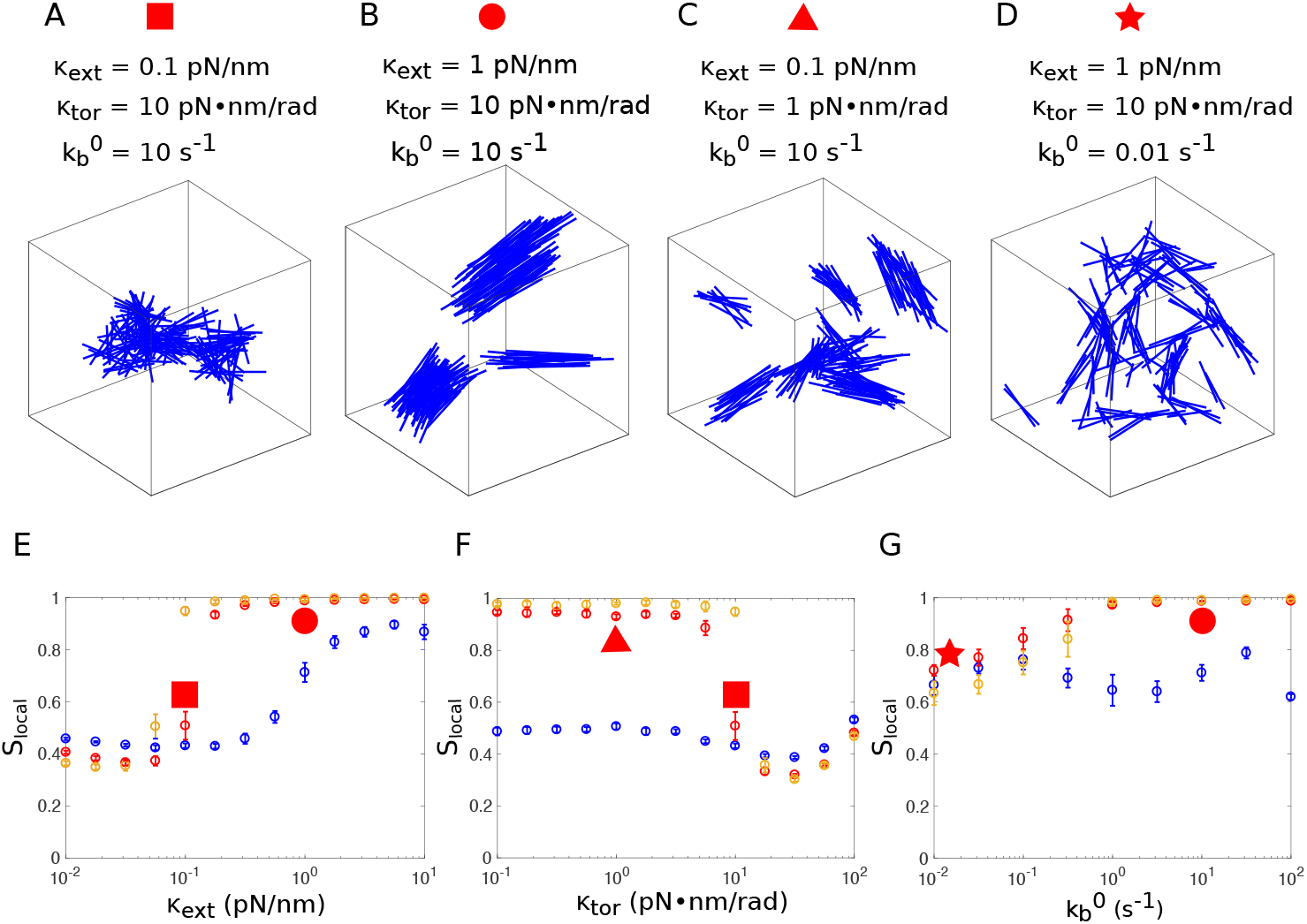
Influence of the crosslinker’s mechanical and kinetic properties on the organization of the actin network. (A-D) Organization of actin filaments of 135nm at the end of the simulation (*t* = 50*s*) for different values of extensional stiffness *κ*_ext_, torsional stiffness *κ*_tor_, and breakage rate 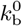, as indicated above the figure. (E) Local nematic order parameter *S*_local_ as a function of the extensional stiffness *κ*_ext_, with *κ*_tor_ = 10pN · nm · rad^−1^ and 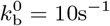. (F) Local nematic order parameter *S*_local_ as a function of the torsional stiffness *κ*_tor_, with *κ*_ext_ = 0.1pN/nm and 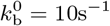. (G) Local nematic order parameter *S*_local_ as a function of the strain-free breakage rate 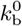, with *κ*_tor_ = 10pN · nm and *κ*_ext_ = 1pN/nm. In (E-G), simulations were performed for filaments of various lengths: 81nm (blue), 135nm (red), 189nm (orange). For each simulation, the means of *S*_local_ were calculated from the data between *t* = 40*s* to 50*s* and the error bars indicate standard deviation over 10 simulations. *S*_local_ corresponding to the networks in panels (A-D) are identified by the red symbols with corresponding shapes.

From the above results, we conclude that crosslinkers with large extensional stiffness favor bundle formation. This result can be intuitively explained by the following simplified but heuristic example involving only two filaments. If two filaments are initially aligned with each other, a slight change in orientation between both filaments results in the stretching of crosslinkers bound at different positions, which leads to a restoring torque to realign the filaments. The torque is proportional to the extensional stiffness and the distance between the positions of attached crosslinkers, thus stiffer crosslinkers create larger realignment torque than softer crosslinkers. Longer filaments not only have more crosslinkers, but also crosslinkers that are more distantly positioned, therefore more extended and more inclined to create a larger torque that will restore the parallel alignment of filaments.

### C. Crosslinkers with high torsional stiffness disfavor bundle formation

We next investigated the impact of torsional stiffness *κ*_tor_ on the organization of actin networks, keeping the extensional stiffness at a relatively small value (*κ*_ext_ = 0.1pN/nm). Torsional stiffness had virtually no influence on the organization of short filaments (81nm) (Fig. 3 F, blue). However, medium length filaments (135nm) had a sharp transition from bundle to meshwork at *κ*_tor_ = 10pN · nm · rad^−1^ (Fig. 3 F, red). A similar trend was also observed for long filaments (189nm) (Fig. 3 F, orange). At highest torsional stiffness tested (*κ*_tor_ = 100pN · nm · rad^−1^), crosslinker attachment lifetime was extremely short because their torsional energy often became much larger than the critical energy *E_c_* that modulates the detachment rate (see Eq. (B1)), therefore the number of attached linkers was significantly reduced (Fig. S2 H [43]). The uncrosslinked actin network formed at this regime was different from the connected meshwork formed at *κ*_tor_ = 10pN · nm · rad^−1^, though their nematic order parameters *S*_local_ were similarly low.

The above results show that crosslinkers with high torsional stiffness disfavor bundle formation. This result can be explained by the fact that, when torsional stiffness is high, formation of several crosslinkers between two aligned filaments results in very high torsional energies, due to the frustrated interaction between crosslinkers. This will become clear later in Sec. III F. Therefore, it is energetically more favorable to form a few but over-stretched crosslinkers with many distant filaments than to form many but under-stretched crosslinkers with a few proximal filaments. In the former situation, filaments form a highly entangled actin meshwork. This explanation is supported by the decreasing number of clusters, *κ*_tor_ (Fig. S2 E [43]), as well as the increasing extensional strains at *κ*_tor_ = 10pN · nm · rad^−1^ (Fig. S4 C [43]).

### D. Turnover of crosslinkers is necessary for large bundle formation

We have shown that high extensional stiffness of crosslinkers favors bundle formation. When the breakage rate 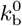 was reduced to 0.01*s*^−1^, even for large extensional stiffness (1pN/nm), filaments formed a structure where small bundles were interconnected but did not align with each other (Fig. 3 D), as indicated by the relatively low local nematic order parameter at 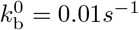 compared with *S*_local_ at 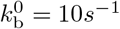 (Fig. 3 G). The difference was even more pronounced in global nematic order parameters *S*_global_ for long filaments (189nm) (Fig. S2 C, orange [43]). At very high breakage rate, the lifetime of bonds between filaments was so short that filaments formed an essentially random, uncrosslinked network. Altogether, we conclude that crosslinker turnover is essential for bundle formation, as alignment of bundles requires the breaking of the bonds that disfavor filament alignment.

### E. Phase diagram of filament organization

Parameter dependence of the actin network structure is summarized in Fig. 4 where we plotted the local nematic order parameter *S*_local_ and the number of attached crosslinkers *N*_attach_ as a function of crosslinking rate constant *k*_f_ and filament length *L*. For a combination of low extensional and high torsional stiffnesses (*κ*_ext_ = 0.1pN/nm, *κ*_tor_ = 10pN · nm · rad^−1^), values of *S*_local_ are concentrated either close to 1 or close to 0.5 (Fig. 4 A, yellow and blue regions, respectively). These two regions are separated by a narrow transition band around *S*_local_ = 0.75 (Fig. 4 A, green). Values of *N*_attach_ = 300 are clustered either close to the saturation number 875, or less than 100, divided by a transition band around *N*_attach_ = 300 (Fig. 4 B, yellow, blue and green regions, respectively). Therefore we chose the lines *S*_local_ = 0.75 and *N*_attach_ = 300 as the boundaries to define the phase diagram (Fig. 4 C). These two lines divide the parameter space into three regions: (1) Above the line *S*_local_ = 0.75, actin filaments are locally aligned into a bundle; (2) Between the lines *S*_local_ = 0.75 and *N*_attach_ = 300, filaments form a crosslinked, disordered meshwork. (3) Below the line *N*_attach_ = 300, filaments are essentially uncrosslinked over the entire simulation time. For larger extensional stiffness (*κ*_ext_ = 1pN/nm), the relative position of the regions is similar, but most of the phase diagram corresponds to bundles (Fig. 4 F), and only the shortest filaments (81nm) form meshworks. In all cases, the existence of three regions in these phase diagrams requires moderate to high breakage rates. When the breakage rate is low 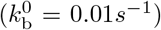, meshworks occupy the entire parameter space (Fig. S3 [43]).

**FIG. 4.**
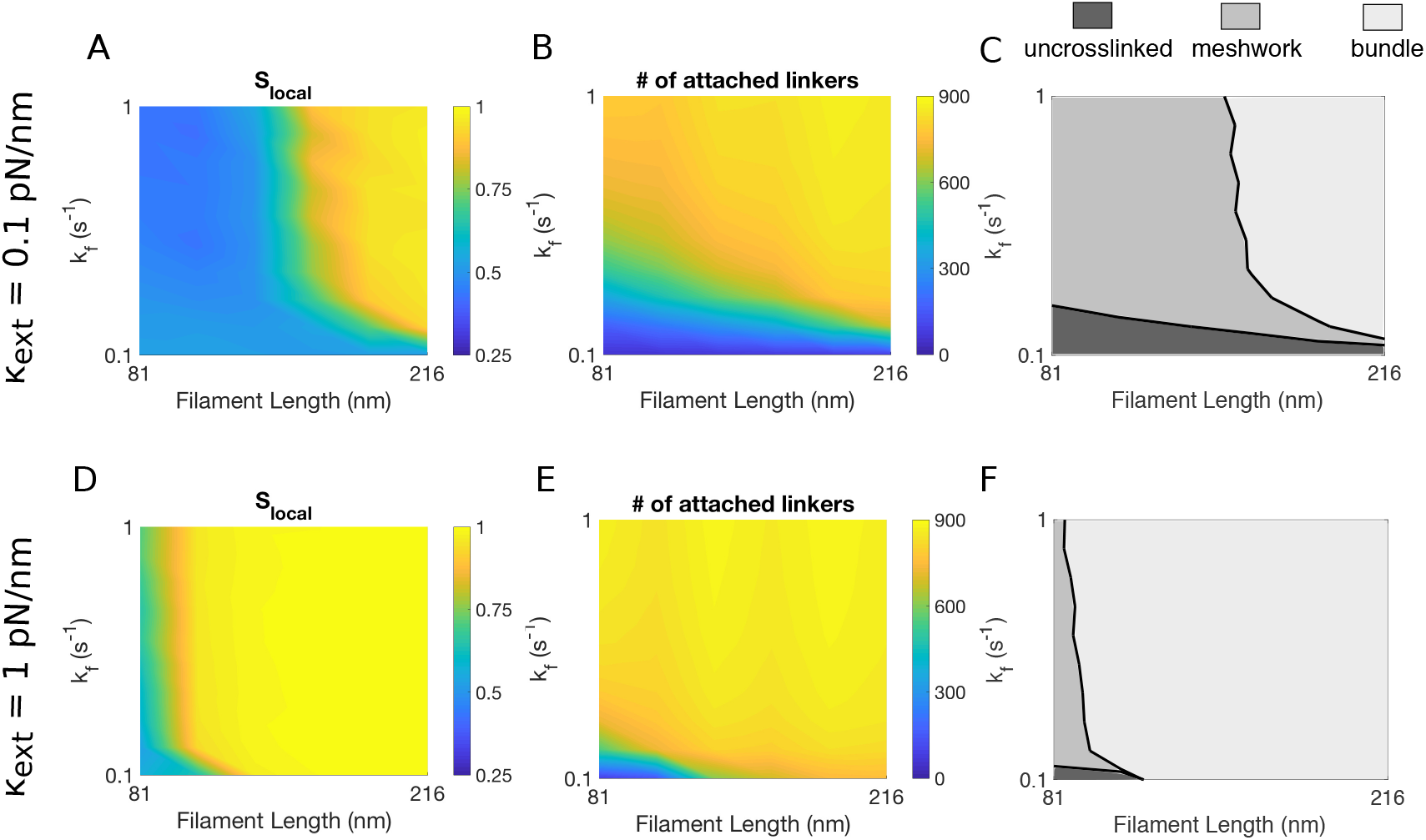
Phase diagram of actin network organization as a function of the crosslinking rate constant *k*_f_ and filament length *L*. (A, D) Local nematic order parameter Slocal as a function of *k*_f_ and *L*. (B, E) Number of attached crosslinkers *N*_attach_ as a function of *k*_f_ and *L*. (C, F) Classification of actin network organizations as a function of *k*_f_ and *L*. The extensional stiffness *κ*_ext_ is 0.1pN/nm in panels (A-C), and 1pN/nm in panels (D-F). In panels (A, B, D, E), plots are constructed by interpolation of results obtained for increment Δ*L* = 27nm of *L* between 81nm and 218nm, and increment Δ*k*_f_ = 0.1*s*^−1^ of *k*_f_ between 0.1*s*^−1^ and 1*s*^−1^. The value for each parameter set is an average over 10 simulations. In (C) and (F), the border separating bundle (light gray) from meshwork (gray) is defined by *S*_local_ − 0.75. The border separating meshwork (gray) from uncrosslinked (dark gray) is defined by *N*_attach_ = 300.

### F. Crosslinked actin networks store elastic energy

Though crosslinkers in our model were not active elements, we found that crosslinkers rapidly became stretched in length, and twisted in angle. To quantify the crosslinkers’ deformation, we introduced the extensional strain *ε* = (*l_c_* − *l*_0_)/*l*_0_, which measures the relative change of a crosslinker’s length *l_c_* from its rest length *l*_0_, and the torsional strain *θ*, which measures the angle between the crosslinker and its bound actin subunits. For all stiffness values, the distributions of the extensional strain *ε* and the torsional strain *θ* significantly deviated from the corresponding Boltzmann distribution for a single independent free spring (Fig. 5 A-C). Strikingly, the extensional strain *ε* peaked at ~ 0.5 but not zero (Fig. 5 A-C, top), indicating that the crosslinkers were stretched on average. The distribution had a narrower width for higher extensional stiffnesses *κ*_ext_. The peak of torsional strain *θ* decreased with increasing torsional stiffness, while the widths of the distributions were essentially the same as in the Boltzmann distribution (Fig. 5 A-C, bottom).

**FIG. 5.**
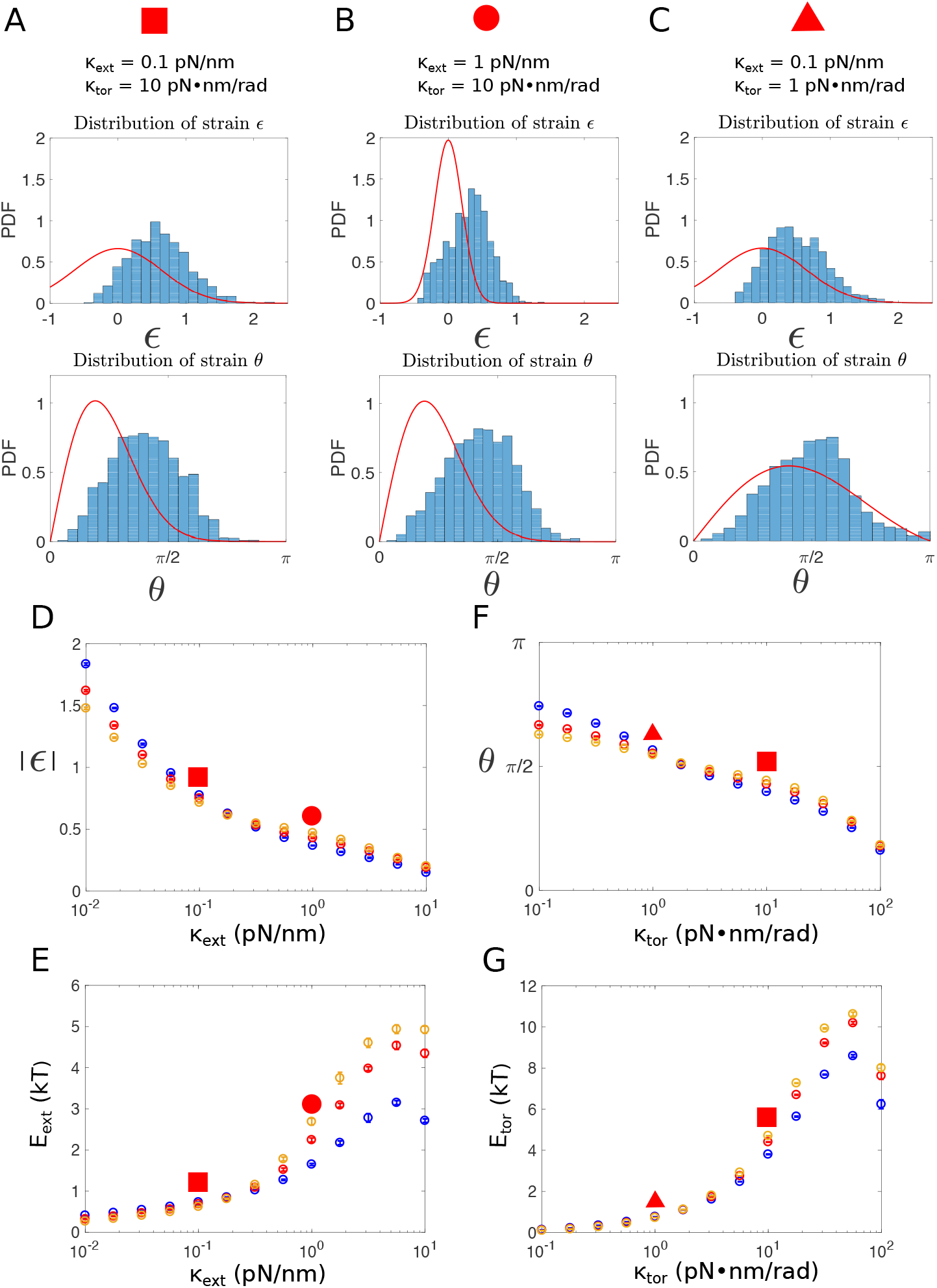
Straining of crosslinkers and energy storage. (A-C) Probability density distribution (PDF) of the extensional strains (top) and torsional strains (bottom) of crosslinkers with various stiffness values as indicated above the figure, at the end of individual simulations (*t* = 50*s*). For comparison, red lines indicate the corresponding Boltzmann distributions of a free spring with the same stiffness. Note that for the extensional strain, when we calculate the Boltzmann distribution, the energy contribution from steric interactions, which lead to the empty region at highly negative strains in the histogram, is neglected. (D, E) Average absolute value of the extensional strain *ε* (D) and the corresponding extensional energy *E_ext_* (E) as a function of the extensional stiffness *κ*_ext_. (F, G) Average torsional strain *θ* (F) and the corresponding torsional energy *E*_tor_ (G) as a function of the torsional stiffness *κ*_tor_. In (D-G), simulations are performed for filaments of various lengths: 81nm (blue), 135nm (red), and 189nm (orange). For each simulation, the means of the energy were calculated from the data between 40s to 50s and the error bars indicate standard deviation over 10 simulations. Energies corresponding to the networks in panels (A-C) are identified by the red symbols with corresponding shapes.

The deviation from the Boltzmann distribution can be primarily accounted by the coupling between crosslinkers that are attached to the same pair of filaments. Indeed, a simpler 1D example of the Brownian motion of two particles each subject to a spring follows similar properties (Fig. S5 [43]). In this example, each spring generates a force of −*κx* if the particle is displaced from the equilibrium position 0 to *x*. If the movement of the two particles were independent (Fig. S5 A [43]), the joint distribution of their positions *x*_1_ and *x*_2_ would simply be the product of identical individual distribution *p*(*x*_1_, *x*_2_) = *g*(*x*_1_)*g*(*x*_2_), where 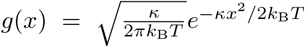 denotes the Boltzmann distribution of an individual particle. However, if the movements of the two particles are coupled, for instance, subject to the constraint *x*_1_ − *x*_2_ = 2*x*_0_ (Fig. S5 B [43]), the position distribution of particle 1 becomes *p*(*x*_1_) = *Cg*(*x*_1_)*g*(*x*_1_ − 2*x*_0_), where C is the normalization constant. Therefore particle 1 is displaced by *x*_0_ due to the coupling with particle 2. Similarly, if two rigid filaments are bound by several crosslinkers, the extensional and torsional strains of these crosslinkers are coupled, and this coupling gives rise to significant strains in the crosslinkers.

We then determined the dependence of the elastic energy stored in the crosslinkers on the crosslinker stiffness. At fixed torsional stiffness *κ*_tor_ = 10pN · nm · rad^−1^, the average magnitude of the extensional strain |*ε*| decreased with increasing extensional stiffness *κ*_ext_ (Fig. 5 D). However, the extensional energy per crosslinker 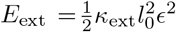 increased (Fig. 5 E), from ~ 0.3*k*_B_*T* for soft extensional springs up to ~ 5*k*_B_*T* for stiff ones, with only a weak dependence on filament length (Fig. 5 E). Similarly, at fixed extensional stiffness *κ*_ext_ = 0.1pN/nm, the average torsional energy per crosslinker *E*_tor_ = *κ*_tor_*θ*^2^ increased with torsional stiffness *κ*_tor_ from ~ 0.2*k*_B_*T* to up to ~ 10*k*_B_*T*, with again a weak dependence on filament length (Fig. 5 G). Both extensional and torsional energies vary relatively smoothly with the corresponding stiffnesses over three orders of magnitude, which is in stark contrast with the sharp structural transition between meshwork and bundle when stiffnesses are varied over the same range (Fig. 3 E and F).

Compared with the increase of the extensional energy *E*_ext_ with *κ*_ext_ (~ 8 − 17 fold), the increase of torsional energy *E*_tor_ with *κ*_tor_ (~ 40 − 50 fold) was more pronounced. This was likely due to stronger coupling between torsional strains of crosslinkers than between extensional strains. Elastic energy plateaued and then slightly decreased for very high stiffnesses (*κ*_ext_ = 10pN/nm or *κ*_tor_ = 100pN · nm · rad^−1^), which was the consequence of higher detachment rate leading to a smaller number of attached linkers, thus reducing the frustrated interactions (Fig. S2 H [43]).

### G. Possible mechanism for torque production by release of the stored elastic energy

We have shown that crosslinking of filaments leads to elastic energy stored in the crosslinkers. How can this energy be transformed into mechanical work? Here, we propose a mechanism for torque generation through orchestrated detachment of crosslinkers by studying a simple model with only two filaments. Let us consider a pair of short filaments, where every other subunit of each filament is crosslinked (Fig. 6). For simplicity, we assume the two filaments are parallel and consider only the rotation of filaments around their axes. We can show that consecutive detachment of crosslinkers from the pointed end to the barbed end lets the filament rotate in the same direction by *π*/13 for each detachment (Appx. F).

**FIG. 6.**
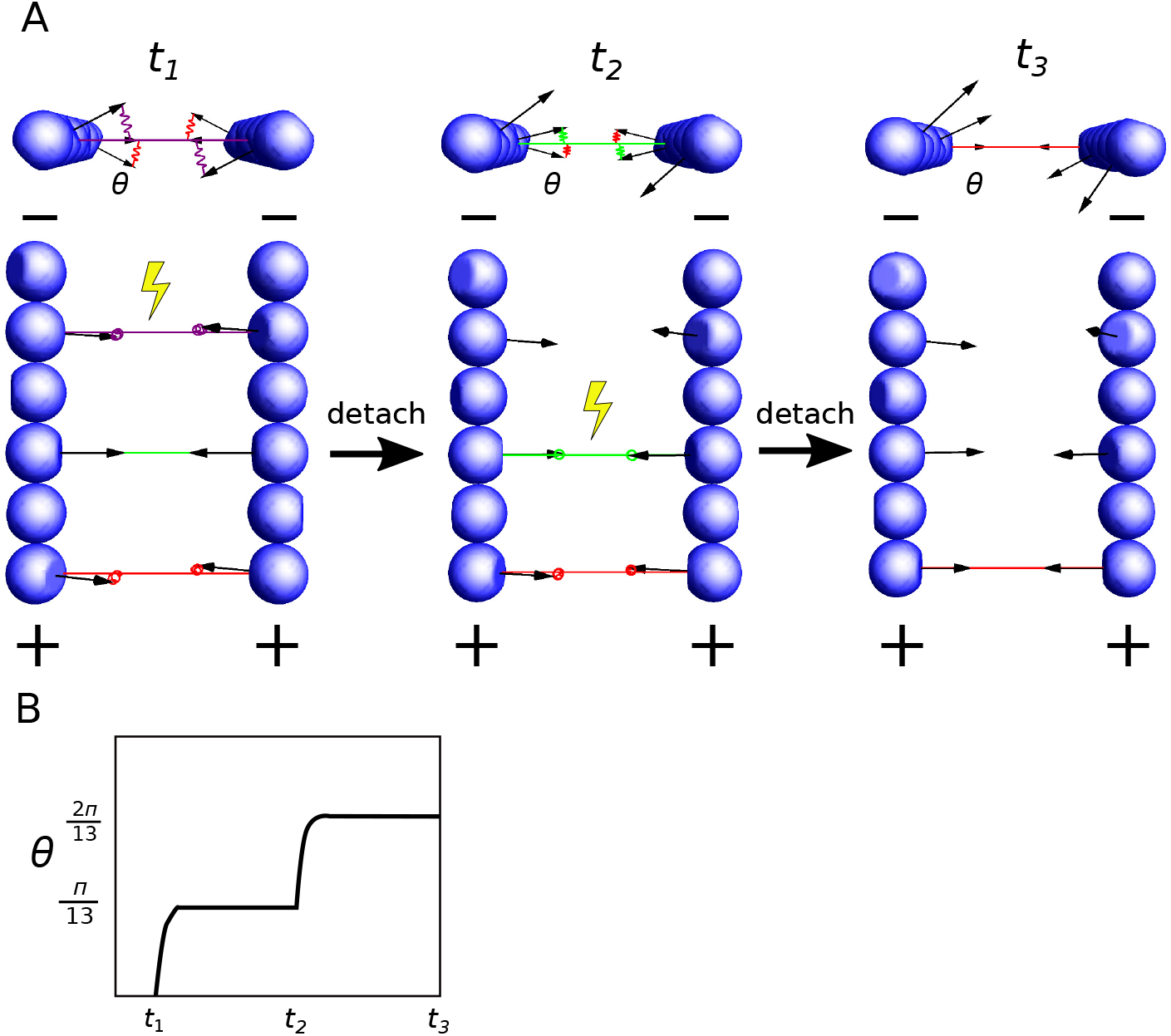
Schematic illustration of a possible mechanism for torque generation by sequential detachment of crosslinkers. (A) Two filaments in parallel are crosslinked on every other subunit. From left to right, crosslinkers are detached from the pointed end (−) to the barbed end (+) sequentially. Upon detachment of a crosslinker (yellow symbols), both filaments rotate around their axes counterclockwise. (B) The angular displacement of the filament upon every detachment is *π*/13.

Building on this simple proof of principle, we can show that sustained directional rotation can be achieved with any filament length and crosslinker spacing and configurations such that (i) the angles between two consecutive crosslinkers along the pair of filaments have the same sign, and (ii) the sum of all these angles is smaller than 2*π* (Appx. F). Under these conditions, breakage of crosslinkers from one end to the other produces directional torque. For example, this ordered crosslinker detachment could be achieved via filaments depolymerization from the pointed end.

## IV. DISCUSSION

### A. Organization of actin filaments in diffraction limited assemblies

In this paper, we showed that highly crosslinked actin networks made of rigid filaments (< 200nm) can form either disordered meshworks or ordered bundles, depending on the filament length and the mechanical and kinetic properties of the crosslinkers. Admittedly, this study does not take into account other actin regulating proteins involved in endocytosis, such as the Arp2/3 complex and capping protein, and thus does not completely resolve the question of the organization of filaments crosslinked with fimbrin at the site of endocytosis in yeast. Even with this limitation, these results provide valuable insights about possible actin filament architectures for endocytosis and other cellular processes that involve short actin filaments. The kinetic properties of fimbrin have been measured *in vitro* [31, 44]. Given the fimbrin concentration present in fission yeast cytoplasm (3.7 *μ*M) [20], these values correspond to rate constants in our model *k*_f_ = 0.2*s*^−1^ and 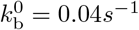. Our simulations suggest that the slow off-rate of fimbrin should favor an assembly of actin filaments into a meshwork (Fig. S3 [43]). Further simulations with branched filaments, and with geometries and dynamics more representative of endocytosis will tell us which type of structure is present at endocytic sites. In addition, further experimental characterization of the mechanical properties of fimbrin and other crosslinkers will be key to understanding the self-organization of actin filaments in diffraction limited structures, and to test the predictions of our simulations.

### B. Mechanisms of energy storage by actin crosslinkers

Our simulations demonstrate that individual actin crosslinkers are able to store up to 10*k*_B_*T* of elastic energy, which is one order of magnitude higher than the elastic energy stored in an uncoupled spring in a thermal bath 1.5*k*_B_*T*, and about half of the energy released by ATP hydrolysis ~ 25*k*_B_*T*. The maximum energy of 10*k*_B_*T* is limited by the “slip-bond” behavior of crosslinkers adopted in our model, i.e., their detachment rate increases with increasing strain (Eq. B1). Some actin binding proteins and crosslinkers demonstrate a “catch-bond” behavior, i.e. their detachment rate decreases with strains in certain range [40, 45, 46]. Crosslinkers with “catch-bond” behavior in principle could store amounts of energy much greater than the values reached in our simulations.

To get a better sense of the amount of energy stored in the crosslinkers, one can make a comparison with the energy necessary to deform the plasma membrane into an endocytic vesicle. A back of the envelope calculation estimates the work needed to create a cylindrical invagination of *R_t_* = 25nm in radius and *D_t_* = 140nm in depth [16] against the turgor pressure *P* ~ 0.8 × 10^6^Pa is 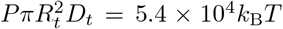. The results of our model suggest that with ~ 900 crosslinkers, they can provide around 10^4^*k*_B_*T* energy, or about 1/6 of the total energy needed. The energy might be even larger if significant turnover of actin takes place during endocytosis, as predicted by mathematical modeling [11].

The main reason energy storage is possible is that short actin filaments are rigid, which creates geometrical constraints on bound crosslinkers, forcing virtually all of them to fluctuate around average lengths and angles that are different from their rest lengths and angles. These frustrated interactions were indeed observed in small-angle X-ray experiments to explain the cooperative binding of actin crosslinkers during actin bundle formation [47].

### C. How can the stored energy be used productively in cellular processes?

So far, only a small fraction of the force necessary to deform plasma membrane during clathrin-mediated endocytosis in yeast can be accounted for by actin polymerization. We predict that at least some of the missing force can come from the conversion of the elastic energy stored in the crosslinkers into force and/or torque.

In this paper, we proposed a specific mechanism for torque production by orchestrated detachment of crosslinkers. This mechanism is different from the Brownian ratchet mechanism of force production that is directly coupled to ATP hydrolysis [48, 49]. However, the ordered detachment has to be coupled to a non-equilibrium process to provide the information necessary for the ordered detachment. Treadmilling of filaments coupled to ATP hydrolysis could play such a role. To estimate the order of magnitude of free energy necessary to provide this information, let us consider a pair of short actin filaments (e.g. 50-subunit long) that are crosslinked by 10 crosslinkers. The free energy cost of detaching the crosslinkers in a specific order among all the 10! possibilities is ~ *k*_B_*T* ln 10! = 13*k*_B_*T*, which only a small fraction of the energy provided by the ATP hydrolysis of two actin filaments undergoing treadmilling (50 × 25 = 1250*k*_B_*T*).

We stress that this ordered detachment is only one possible mechanism to use the energy and more mechanisms need to be discovered. Future work with more realistic models for endocytosis or other actin-based processes will likely uncover new orchestrated mechanisms for force production. We also predict that the energy stored in the crosslinked meshwork can be converted into force and torque by stochastic breakages in the stressed meshwork (similar to snapping a pre-tensed rubber band, for example), and along the same lines as the mechanisms that have been proposed for symmetry breaking and force production in actin-based motility systems [50–52].

## Appendix A: Model of actin filaments

Actin filaments are modeled as rigid cylindrical rods with diameter *b* and length *L*. The position of a filament is represented by its center of mass **C**. A unit vector **N** pointing from the filament’s pointed end to the barbed end indicates the orientation of the filament. The *i*-th subunit (counting from the pointed end) carries a unit vector **O**_*i*_, which is normal to the binding surface with a crosslinker (Fig. 1A). We assume all **O**_*i*_-s are perpendicular to the filament’s orientation **N**. Two consecutive subunits **O**_*i*_ and **O**_*i*+1_ span an angle of 14*π*/13 calculated counter-clockwise from **O**_*i*_ to **O**_*i*+1_. This mean two consecutive subunits on different strands have their binding interface in almost opposite directions, and two consecutive subunits on the same strand have their binding interface at an angle of 2*π*/13 ( 28°). We arbitrarily choose the filament’s rotational vector M as the normal vector of the first subunit **M** = **O**_1_. The orientational degree of freedom of the filament thus is fully captured by three orthonormal vectors **N, M** and **N** × **M**.

The motion of a filament is described by its translational velocity **V**_*c*_ and angular velocity **Ω**, which are defined by the following equations:

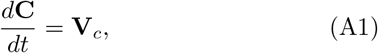

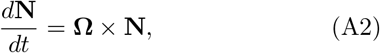

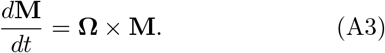

The velocities **V**_*c*_ and **Ω** are governed by the force-balance and torque-balance equations:

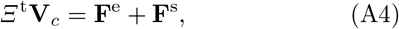

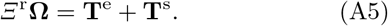

Here the 3 × 3 matrices *Ξ*^t^ and *Ξ*^r^ denote the frictional matrix associated with translational and rotational motion of the filament, respectively. The vectors **F**^e^ and **T**^e^ denote the total deterministic force and torque generated by crosslinkers or induced by steric interactions between filaments. The vectors **F**^s^ and **T**^s^ denote the stochastic force and torque, which obey the fluctuation-dissipation relations:

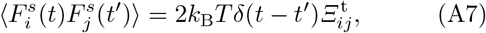

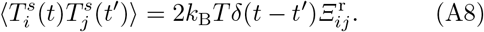

Here the subscript indicates the element of the vectors or matrices. The frictional matrices are anisotropic, and given by [53]:

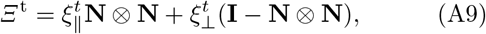

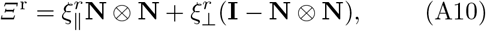

where 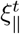 and 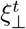 are the frictional coefficients for translational movement parallel with and perpendicular to the filament’s central axis, and 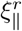 and 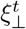 are the corresponding frictional coefficients for rotation. The 3 × 3 identity matrix is denoted by **I**, and ⊗ denotes the outer product of two vectors. The anisotropic frictional coefficients depend on filament length *L* and diameter *b* via the relations [54]:

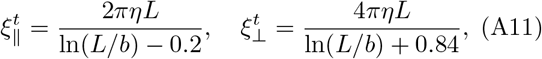

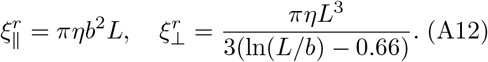

Here *η* denotes the viscosity of the medium.

To account for the steric interaction between filaments, if the shortest distance *r*_min_ between two filaments is less than the diameter *b* of a filament, a constant repulsive force *f*_st_ is applied along the lines connecting the two nearest points.

In each time step, we calculate all the forces and torques acting on a filament and determine the translational velocity **V**_c_ and angular velocity **Ω** of the filament according to Eqs. (A4) and (A5). The center of mass of a filament is then updated as:

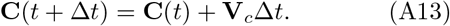

The updated orientations are:

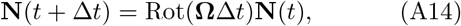

where Rot(**Ω**Δ*t*) denotes the rotation matrix defined by the vector **Ω**Δ*t*. The rotation vector **M**(*t*) is updated in the same way.

## Appendix B: Model of actin crosslinking proteins

Each actin crosslinking protein is modeled as an elastic spring that bridges two actin subunits in two separate filaments. The crosslinking of two unoccupied subunits proceeds with a rate constant of *k*_f_, as long as the subunits are less than *r*_c_ apart. The breakage of an established crosslinker is assumed to follow a “slip-bond” mechanism and occurs with an energy-dependent rate constant:

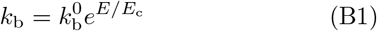

where 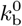 denotes the strain-free breakage rate constant, *E* denotes the total elastic energy, and *E_c_* denotes the critical energy that determines the sensitivity of the bond breakage on the forces and torques.

The elastic energy *E* of a crosslinker that bridges actin subunits in filaments *α* and *β* is a function of the positions, orientations and rotations of both filaments, *E* = *E*(**C**^*α*/*β*^, **N**^*α*/*β*^, **M**^*α*/*β*^). The force generated by the crosslinker on filament *α* reads:

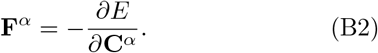

To determine the torque generated by the crosslinker on filament *α*, we choose three orthnormal vectors **e**_1_, **e**_2_, **e**_3_ and virtually rotate filament *α* by an infinitesimal angle *φ_i_* around the axis **e**_*i*_. These operations are equivalent to applying the following infinitesimal changes to the orientational vectors of filament:

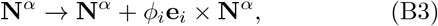

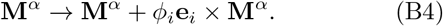

The elastic energy correspondingly has an infinitesimal change *E* → *E* + Δ*E*. The torque then reads:

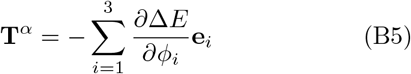

Forces and torques acting on filament *β* are derived in a similar way. The total elastic force and torque are obtained by summing (B2) and (B5) over all the crosslinkers bound to the filament.

## Appendix C: Events performed in a single simulation step

At each time step Δ*t* of the simulation, we perform the following operations:

1. Calculate all the deterministic and stochastic forces and torques on each filament. These forces and torques determine the translational and angular velocities of filaments according to Eqs. (A4) and (A5). The positions and orientations of filaments are then updated according to Eqs. (A13) and (A14).
2. For each pair of filament *α* and *β*, determine the number *N_a_* of crosslinkers to be attached between them by drawing a random number from the Poisson distribution, Pois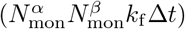, where 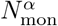 and 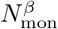 are the number of actin subunits in each filament. Then randomly pick *N_a_* pairs of subunits (*i, j*) with *i* in filament *α* and *j* in filament *β*. If they are not already occupied by a crosslinker and distance between them is shorter than the interacting distance *r_c_* and the total number of already occupied subunits in each filament is less than 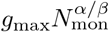, a crosslinker is built between the subunits (*i, j*).
3. For each already existing crosslinker, determine whether it will be detached by comparing a uniformly distributed random number *u* on the interval [0, 1] with the energy-dependent breakage rate *k*_b_ in Eq. (B1). If *u* < 1 − *e*^−*k*_b_Δ*t*^, the crosslinker is detached.

In our simulation, we always set the time step Δ*t* at least 100 times smaller than the relaxation time of the spring 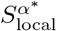 to ensure that we correctly capture the dynamics of the springs. For computational reasons, we used a high viscosity value *η* = 10Pa · s, such that the relaxation time *τ* ~ 0.01s for *κ*_ext_ = 0.1pN/nm and *κ*_tor_ = 10pN · nm · rad^−1^. We tested values of lower viscosity down to *η* = 0.1Pa · s. There is no significant difference in the local nematic order parameter *S*_local_ and the elastic energies *E*_ext_ and *E*_tor_ between *η* = 0.1Pa · s and *η* = 10Pa · s. However, the global nematic order parameter *S*_global_ for long filaments is increased to 1 at lower viscosity (Fig. S7 [43]). This is because the enhanced diffusion increases the probability of filaments moving close to each other. As a result, the separated bundles observed at high viscosity merge into a single bundle when the viscosity is low, increasing the global nematic order parameter.

## Appendix D: Metrics to describe local and global organization of filaments

We characterize the structure of actin clusters by introducing local and global nematic order parameter *S*_local_ and *S*_global_. We map the connections between filaments into an undirected graph, with filaments being the nodes, and the number of crosslinkers being the value at the edges connecting two nodes. Filaments in a connected component of the graph are said to form a cluster if the number of filaments in the component is more than 10. The nematic order parameter *S* for a group of filaments is the maximum eigenvalue of the following matrix [55]:

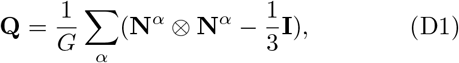

where *G* denotes the number of filaments in the group, **N**^*α*^ denotes the orientational vector of filament *α*. For global nematic order parameter *S*_global_, the group in Eq. (D1) includes all the filaments. For a particular filament *α*\*, 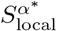 is defined by grouping the filament *α*\* and its connected nodes in Eq.(D1). The local nematic order parameter *S*_local_ is the average of 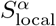 over all the filaments that have at least 2 connected nodes.

Both *S*_local_ and *S*_global_ are in the range of [0, 1]. Values of *S*_local_ close to 1 indicate that filaments are locally aligned with their connected neighbors. Values of Sglobal close to 1 indicate that all the filaments are aligned. In general *S*_local_ is greater than *S*_global_, and reaches steady state more rapidly, because filaments that are in close proximity can rapidly aligned, but it takes time for distant clusters of filaments to collide and reorient. *S*_local_ is also more consistent over different simulations than Sglobal, as reported by smaller error bars for Slocal than for *S*_global_ (e.g. Fig. 2G). If the viscosity of the medium *η* is reduced to 1Pa · s, filaments form a single cluster and the error bars of *S*_global_ become comparable with *S*_local_ (Fig. S7 B,C [43]).

Note that in a sparsely connected network with filaments in random orientation, *S*_local_ ~ 0.5 (Fig. S1 B [43]) is higher than one should expect ~ 0. This artifact is due to the fact that the sum in Eq. (D1) is done over a very small number of filaments (~ 3). We confirmed this property by numerically calculating the nematic order parameter for three unit vectors with random orientations. The resulting distribution of Slocal has a peak at 0.45 (Fig. S6 [43]). This almost uncrosslinked network should be distinguished from the densely connected actin meshworks, which have local nematic order parameters *S*_local_ in the same range (~ 0.4 Fig. 2 B) but possess a large number of attached crosslinkers. Therefore, the number of crosslinkers in the meshwork is required to distinguish these two structures.

## Appendix E: Validity of the rigidity assumption

In our model, we assumed that filaments are rigid. This rigidity assumption implies: (i) thermal fluctuations do not significantly bend or twist the filament; (ii) forces and torques exerted by crosslinkers do not significantly bend or twist the filaments. We can verify a posteriori that these conditions are actually fullfiled in our simulations. Indeed, the maximum force produced by a crosslinker in our simulations is ~ 10pN when the extensional stiffness *κ*_ext_ reaches 10pN/nm, and the maximum torque is ~ 100pN · nm when the torsional stiffness *κ*_tor_ reaches 100pN · nm · rad^−1^. Given the persistence length of actin filament for both bending and twisting is *L_p_* ~ 10*μ*m [12, 14, 56, 57], for a filament of length *L* = 200nm which consists of *N* = *L*/*δ* = 74 subunits, the angular change between two consecutive subunits due to thermal fluctuation is arccos(*e*^−*L*/*L_p_*^)/*N* = 0.15°. The angular change due to bending caused by a force of *f* = 10pN in the middle of the filament, while the two ends are fixed, is arctan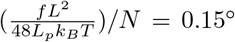 [58]. The twisting angle by a torque of *T* = 100pN · nm is 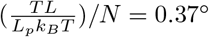. Therefore, it is safe to consider filaments as stiff, and the energy stored in crosslinkers would not be dramatically different even if the finite stiffness of filaments was taken into account.

## Appendix F: Direct rotation of filaments by consecutive detachment of crosslinkers

We consider the rotation of two parallel filaments around their axes by consecutive detachment of crosslinkers from the pointed end to the barbed end. We assume that every other subunit of each filament is crosslinked, such that the *i*-th crosslinker has an angle of

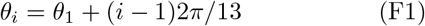

with its attached actin subunit. The torque generated by the *i*-th crosslinker on the filament thus is −*κ*_tor_*θ_i_*. Here the crosslinker label *i* is ordered according to their distance to the pointed end of filaments. At torque balanced state, i.e., 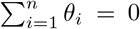, we have *θ*_1_ = −(*n* − 1)*π*/13. Upon detachment of crosslinker 1, the total torque becomes imbalanced and the filament makes a rotation of angle Δ*φ* to reach a new torque balanced state, i.e., 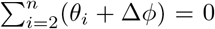. This leads to Δ*φ* = −*π*/13. Similarly we can show that attachment of a new crosslinker at the (*n* + 1)-th position *θ*_*n*+1_ = *θ*_1_ + *n*2*π*/13 will cause the filament rotate the same angle in the same direction as caused by detachment of the 1-st crosslinker.

Note that even though the rotation angle Δ*φ* is independent of the number of attached crosslinkers, it is required that crosslinkers are present in large enough number or are stiff enough to ensure that rotation will be significantly larger than thermal fluctuations, i.e. 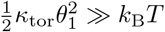. For instance, if the number of crosslinkers n = 10, this requires *κ*_tor_ ≫ 2pN · nm · rad^−1^.

The above calculation can be easily extended to situations where Eqs. (F1) do not hold. The angular rotation Δ*φ*^(*i*)^ of the filament upon detachment of the *i*-th crosslinker satisfies the recursive relation:

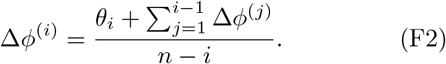

Directed rotation requires that Δ*φ*^(*i*)^ have the same sign for all *i*. This is equivalent to that the angles between consecutive crosslinkers (*θ_i_* − *θ*_*i*−1_) have the same sign for all *i*.

## ACKNOWLEDGMENTS

We thank the members of the Berro lab for helpful discussions and Prof. Jonathan Howard, Prof. Enrique De La Cruz, Prof. Michael Murrel, Dr. Wenxiang Cao, Dr. Pablo Sartori, Danny Seara for critical reading of the manuscripts. This research was supported in part by National Institutes of Health/National Institute of General Medical Sciences Grant R01GM115636.

## References

[1] D. C. Amberg, Mol Biol Cell 9, 3259 (1998).

[2] D. R. Kovar, V. Sirotkin, and M. Lord, Trends Cell Biol 21, 177 (2011).

[3] E. Bi and H. O. Park, Genetics 191, 347 (2012).

[4] J. B. Moseley and B. L. Goode, Microbiol Mol Biol Rev 70, 605 (2006).

[5] T. D. Pollard and J. Q. Wu, Nat Rev Mol Cell Biol 11, 149 (2010).

[6] H. Tang, T. C. Bidone, and D. Vavylonis, Cytoskeleton (Hoboken) 72, 517 (2015).

[7] T. C. Bidone, H. Tang, and D. Vavylonis, Biophysical journal 107, 2618 (2014).

[8] H. Tang, D. Laporte, and D. Vavylonis, Molecular biology of the cell 25, 3006 (2014).

[9] M. R. Stachowiak, C. Laplante, H. F. Chin, B. Guirao, E. Karatekin, T. D. Pollard, and B. O’Shaughnessy, Dev Cell 29, 547 (2014).

[10] A. Michelot, A. Grassart, V. Okreglak, M. Costanzo, C. Boone, and D. G. Drubin, Dev Cell 24, 182 (2013).

[11] J. Berro, V. Sirotkin, and T. D. Pollard, Mol Biol Cell 21, 2905 (2010).

[12] F. Gittes, B. Mickey, J. Nettleton, and J. Howard, J Cell Biol 120, 923 (1993).

[13] H. Isambert, P. Venier, A. C. Maggs, A. Fattoum, R. Kassab, D. Pantaloni, and M. F. Carlier, J Biol Chem 270, 11437 (1995).

[14] J. S. Graham, B. R. McCullough, H. Kang, W. A. Elam, W. Cao, and E. M. De La Cruz, PLoS One 9, e94766 (2014).

[15] J. Berro, A. Michelot, L. Blanchoin, D. R. Kovar, and J. L. Martiel, Biophys J 92, 2546 (2007).

[16] W. Kukulski, M. Schorb, M. Kaksonen, and J. A. Briggs, Cell 150, 508 (2012).

[17] B. J. Galletta and J. A. Cooper, Curr Opin Cell Biol 21, 20 (2009).

[18] O. L. Mooren, B. J. Galletta, and J. A. Cooper, Annu Rev Biochem 81, 661 (2012).

[19] M. Kaksonen, Y. Sun, and D. G. Drubin, Cell 115, 475 (2003).

[20] V. Sirotkin, J. Berro, K. Macmillan, L. Zhao, and T. D. Pollard, Mol Biol Cell 21, 2894 (2010).

[21] A. Picco, M. Mund, J. Ries, F. Nedelec, and M. Kaksonen, Elife 4 (2015), 10.7554/eLife.04535.

[22] I. Gaidarov, F. Santini, R. A. Warren, and J. H. Keen, Nat Cell Biol 1, 1 (1999).

[23] C. J. Merrifield, M. E. Feldman, L. Wan, and W. Almers, Nat Cell Biol 4, 691 (2002).

[24] M. Kaksonen, C. P. Toret, and D. G. Drubin, Cell 123, 305 (2005).

[25] D. Loerke, M. Mettlen, S. L. Schmid, and G. Danuser, Traffic 12, 815 (2011).

[26] E. Cocucci, F. Aguet, S. Boulant, and T. Kirchhausen, Cell 150, 495 (2012).

[27] Y. Tseng, T. P. Kole, J. S. Lee, E. Fedorov, S. C. Almo, B. W. Schafer, and D. Wirtz, Biochemical and biophysical research communications 334, 183 (2005).

[28] T. Kim, W. Hwang, H. Lee, and R. D. Kamm, PLoS computational biology 5, e1000439 (2009).

[29] T. Kim, W. Hwang, and R. D. Kamm, Biophysical journal 101, 1597 (2011).

[30] E. Kubler and H. Riezman, EMBO J 12, 2855 (1993).

[31] C. T. Skau, D. S. Courson, A. J. Bestul, J. D. Winkelman, R. S. Rock, V. Sirotkin, and D. R. Kovar, Journal of Biological Chemistry 286, 26964 (2011).

[32] N. Minc, A. Boudaoud, and F. Chang, Curr Biol 19, 1096 (2009).

[33] B. Goldenbogen, W. Giese, M. Hemmen, J. Uhlendorf, A. Herrmann, and E. Klipp, Open Biol 6 (2016), 1098/rsob.160136.

[34] R. Basu, E. L. Munteanu, and F. Chang, Mol Biol Cell 25, 679 (2014).

[35] S. Dmitrieff and F. Nedelec, PLoS Comput Biol 11, e1004538 (2015).

[36] D. J. Tweten, P. V. Bayly, and A. E. Carlsson, Phys Rev E 95, 052414 (2017).

[37] M. J. Footer, J. W. Kerssemakers, J. A. Theriot, and M. Dogterom, Proc Natl Acad Sci U S A 104, 2181 (2007).

[38] S. Pyrpassopoulos, G. Arpag, E. A. Feeser, H. Shuman, E. Tuzel, and E. M. Ostap, Sci Rep 6, 25524 (2016).

[39] M. J. Greenberg, G. Arpag, E. Tuzel, and E. M. Ostap, Biophys J 110, 2568 (2016).

[40] J. M. Laakso, J. H. Lewis, H. Shuman, and E. M. Ostap, Science 321, 133 (2008).

[41] M. M. Claessens, M. Bathe, E. Frey, and A. R. Bausch, Nat Mater 5, 748 (2006).

[42] X. J. Janssen, J. M. van Noorloos, A. Jacob, L. J. van Ijzendoorn, A. M. de Jong, and M. W. Prins, Biophys J 100, 2262 (2011).

[43] See Supplemental Material at [] for details.

[44] C. T. Skau and D. R. Kovar, Curr Biol 20, 1415 (2010).

[45] D. L. Huang, N. A. Bax, C. D. Buckley, W. I. Weis, and A. R. Dunn, Science 357, 703 (2017).

[46] T. Z. Luo, K. Mohan, P. A. Iglesias, and D. N. Robinson, Nature Materials 12, 1063 (2013).

[47] H. Shin, K. R. P. Drew, J. R. Bartles, G. C. Wong, and G. M. Grason, Physical review letters 103, 238102 (2009).

[48] A. Mogilner and G. Oster, Biophysical journal 84, 1591 (2003).

[49] A. Mogilner and G. Oster, Biophysical journal 71, 3030 (1996).

[50] K. Sekimoto, J. Prost, F. Julicher, H. Boukellal, and A. Bernheim-Grosswasser, Eur Phys J E Soft Matter 13, 247 (2004).

[51] A. Kawska, K. Carvalho, J. Manzi, R. Boujemaa-Paterski, L. Blanchoin, J. L. Martiel, and C. Sykes, Proceedings of the National Academy of Sciences of the United States of America 109, 14440 (2012).

[52] K. Carvalho, J. Lemiere, F. Faqir, J. Manzi, L. Blanchoin, J. Plastino, T. Betz, and C. Sykes, Philos Trans R Soc Lond B Biol Sci 368, 20130005 (2013).

[53] C. H. Schreiber, M. Stewart, and T. Duke, Proc Natl Acad Sci U S A 107, 9141 (2010).

[54] J. Howard, Mechanics of motor proteins and the cytoskeleton (Sinauer Associates, Publishers, Sunderland, Mass., 2001) pp. xvi, 367 p.

[55] M. Doi, Soft matter physics (Oxford University Press, 2013).

[56] R. Yasuda, H. Miyata, and J. Kinosita, K., J Mol Biol 263, 227 (1996).

[57] Y. Tsuda, H. Yasutake, A. Ishijima, and T. Yanagida, Proc Natl Acad Sci U S A 93, 12937 (1996).

[58] J. M. Gere and S. P. Timoshenko, PWS-KENT Publishing Company 534, 4 (1997).

